# Phenotypic plasticity meets and moulds carry-over effects in sea rock-pool mosquitoes

**DOI:** 10.1101/2024.07.24.604954

**Authors:** Giulia Cordeschi, Roberta Bisconti, Valentina Mastrantonio, Daniele Canestrelli, Daniele Porretta

## Abstract

Environmental conditions experienced during early-life can shape trait expression after metamorphosis. Direct carry-over effects occur when the value of a trait expressed at an early stage directly determines its expression at later stages, maintaining phenotypic continuity across metamorphosis. However, environmental factors generally affect multiple traits whose interaction might shape developmental trajectories. Here, we tested the hypothesis that trait interactions can shape direct carry-over effects by examining the interplay between behavioural and morphological plasticity in the sea rock-pool mosquito *Aedes mariae* under varying salinity conditions. We found plastic changes in body size and behaviour. Higher salinity reduced larval body size and increased resting behaviour at the water surface while browsing activity decreased. Under constant conditions, larval body size was positively correlated with pupal size, indicating a direct carry-over effect. However, this relationship was disrupted as salinity increased due to larval size-dependent behavioural plasticity which decouples pupal morphology from larval size. Our results show that environmental conditions modulate trait integration and modify direct carry-over effects. These findings highlight the importance of considering multiple traits when studying developmental plasticity and contribute to the ongoing debate on the extent to which one life stage is coupled to the others across metamorphic boundary.

## Introduction

The environment varies across space and time, and phenotypic plasticity is one of the major mechanisms enabling organisms to respond to such variability [1–3]. Phenotypic plasticity is the ability of a given genotype to respond to environmental change by expressing different phenotypes [4,5]. In animals undergoing metamorphosis, environmental conditions experienced early in life can shape trait expression across this event of major change [4,6–10]. This phenomenon, known as the carry-over effect, is a form of developmental plasticity [7–9,11,12]. When environmental factors influence the expression of a trait in an earlier life stage and this effect is directly transferred to subsequent stages, it is referred to as a ‘direct carry-over effect’ [8]. Typical cases of direct carry-over effects are size at metamorphosis in marine invertebrates, insects and amphibians, where larval body size determines post-metamorphic body size [13–15]. For example, in wood frogs (*Rana sylvatica*), canopy cover affects larval size, with individuals from closed-canopy ponds growing larger than those from open-canopy ponds—an effect that persists after metamorphosis [14,16].

Nevertheless, more complex patterns have recently emerged [17–21]. Indeed, organisms often respond to environmental change with plasticity in multiple traits (i.e. multivariate plasticity) [22,23]. Well-documented examples include predator-induced defences in aquatic animals, where prey modify behavioural, morphological, and life-history traits in response to predator presence [24–26]. Moreover, plastic responses in one trait can, in turn, shape the development and selection of other traits (reviewed in 27). For instance, the caterpillar *Battus philenor* responds to high temperatures by changing its body colour from black to red and leaving host plants to seek cooler locations for thermal refuge [27]. Experimental studies have shown that red colouration reduces the frequency of refuge-seeking, suggesting that the interaction between these two plastic traits is mediated by the effect of colour on body temperature [28]. This multivariate perspective has provided valuable insights into the trade-offs between the costs and benefits of plasticity, shedding light on evolutionary constraints, adaptation potential, and how populations persist in rapidly changing or novel environments [29–33]. However, when considering plasticity across metamorphic boundaries, the extent to which interactions between multiple plastic traits in early life stages influence the magnitude of carry-over effects remains unclear.

Empirical evidence supports the idea that plastic changes in behaviour can influence individual morphological or physiological traits [31,34–37], Here, we tested the hypothesis that such trait interactions can shape direct carry-over effects. Specifically, we investigated whether plasticity in behavioural traits interferes with the direct carry-over of body size across metamorphosis in the sea rock-pool mosquito *Aedes mariae*. This species undergoes four distinct life stages (egg, larva, pupa, and adult) [38] and inhabits sea-rock pools of supralittoral zone along the western Mediterranean coast [39,40]. Rock pools are extremely variable habitats, experiencing high salinity fluctuations from freshwater to hypersaline conditions over extremely short time frames [41,42]. Using an individual-based approach, we tracked larval and pupal development under two different salinity conditions. We measured larval activity and body size throughout metamorphosis to assess plasticity in these traits and performed path analysis to infer causal relationships between them during development.

## Material and Methods

### Sampling and experimental conditions

Experiments were carried out on individuals of *Ae. mariae* obtained from eggs collected in July 2020 from supralittoral rock pools of San Felice Circeo, Italy (41°13’18.77 “N, 13° 4’5.51”E). We designed two experimental treatments based on field data collected during the reproductive season of the species. In the first treatment, individuals were maintained at a constant 50 ‰ salinity (50 g/l) throughout the experiment (constant condition, hereafter C). In the second treatment, larvae were exposed to increasing salinity from 50 ‰ to 150 ‰, with an increase of 10 ‰ per day throughout the experiment (salinity condition, hereafter S). The experiments were run in a climate chamber set to 26 ± 1 °C, 14 h light and 10 h dark regime. Experimental larvae were fed daily with 1 mg of cat food [43].

### Morphological and behavioural trait measurements

Second instar larvae were individually placed into plastic trays (12x12x7 cm) filled with 200 ml of tap water, previously salted with aquarium salt (Tetra Marine Seasalt). Then, we randomly assigned individuals to constant or salinity developmental treatments (N = 30 x treatment). We placed larvae individually to follow the development of each individual across its life cycle from larval stages to pupae.

We measured the fourth instar larvae’s body size and spontaneous activity. Morphometric measures were obtained by taking digital pictures using a stereomicroscope Leica EZ4W at magnification 1x of all individuals within two hours from the ecdysis. Subsequently, using the open-access software IMAGEJ we measured the width and length of the head, thorax, abdomen, and total body [44]. Larval spontaneous activity was analysed by video-recording larvae for 10 min in their housing containers using a camera Canon TG-6. Individuals were tested within 12 hours from the ecdysis between 8 am and 1 pm. A single operator (G.C.), blind to the treatment, manually scored videos using the software BORIS (Behavioral Observation Research Interactive Software). Larval behaviours in the ethogram consisted of resting and browsing. During resting behaviour, the larva is positioned underwater or at the water surface with the respiratory siphon attached to the air-water interface and the body hanging obliquely into the water column. In browsing behaviour, the larva brushes the wall or the bottom of the container with its mouthparts and moves along the container surface [45].

For the pupal stage, we measured the pupal body size. Measurements were obtained from digital images, following the same procedure described for the larval stage. We measured cephalo-thorax width [44,46] and determined the sex of individuals by checking the last cephalo-thorax segment [38].

### Data analysis

To find evidence of plastic response in larval morphology and spontaneous activity to salinity conditions, we carried out Generalised linear models (GLMs). We applied principal component analysis (PCA) on fourth-instar larvae morphometric measures and subsequently performed GLM on the first two principal components separately as dependent variables. The analysis was run using the *glm* function in R with a Gaussian distribution and with identity link function for PCA components of larval morphology and larval resting behaviour. We applied GLMs with Gamma distribution and inverse link for browsing behaviour. We relied on the Akaike information criterion (AIC) to identify the distribution type and the link function that improved the model fitting. We checked for the effect of sex because of the sexual dimorphism in this mosquito species, although it is usually not evident at the larval stage [47]; however, for models where there was no evidence for an effect of sex based on likelihood ratio test (LRT), we did not include this term in the final models (see Table S2) .

To infer causal relationships between morphological and behavioural traits along the development, we performed path analysis using piecewise structural equation approach using *piecewiseSEM* package. Structural equation models (SEM) provide means to link multiple predictor and response variables in a single casual network, in which paths indicate hypothesised relationships between variables [48]. Structural equations modelling is especially useful when response variables can also act as predictor variables of another response variable—that is, when they mediate an indirect relationship. Since we hypothesised that the body size of early developmental stages might determine the body size of the successive stage and expected an influence of the activity behaviour in shaping body size throughout the development, we included three linear relationships in our path analysis (lm models). We fit two separate SEM for treatments since we hypothesised that the covariation patterns depend on water salinity. We used in the model only resting behaviour because of the significantly high correlation between resting and browsing measures (r = -0.91, *P*<0.001). The coefficients were standardised to allow comparison between variables.

Using the *multigroup* function, we performed a multigroup analysis to evaluate differences in path coefficients between salinity conditions models. This analysis implements a model-wide interaction in which every term in the model interacts with the grouping variable (i.e., constant versus increasing salinity). If the interaction is significant, the path is different between salinity conditions and is free to vary by group; if not, the path is constrained and takes on the estimate from the global dataset.

All statistical analyses were performed in R version 4.4.2.

## Results

### Plastic response of morphological and behavioural larval traits

The first PCA component explained 76.9% of the total variance and discriminated against individuals on body size. The second component explained 13.9 % of the variance and detected differences in body shape (Table S1).

The GLM conducted on larval body size (PC1) showed significant differences between treatments (Table 1). Fourth instar larvae grown up in increasing salinity treatment were smaller than larvae in constant treatment (Fig. 1A). We found no effect of treatment on the larval shape (PC2; Table 1, Fig. 1B).

**Table 1.** Outcome of generalised linear models using treatment as fixed factor. Significant *P*-values are shown in bold.

| Trait | Coefficient | Standard error | $t$-value | $P$-value |
| --- | --- | --- | --- | --- |
| Larval body size- PC1 | -2.583 | 0.532 | -4.849 | **< 0.001** |
| Larval body shape - PC2 | -0.460 | 0.266 | -1.728 | 0.090 |
| Larval spontaneous activity-Resting behaviour | 0.077 | 0.038 | 2.02 | **0.048** |
| Larval spontaneous activity-Browsing behaviour | 9.876 | 3.914 | 2.523 | **0.014** |

**Figure 1.**
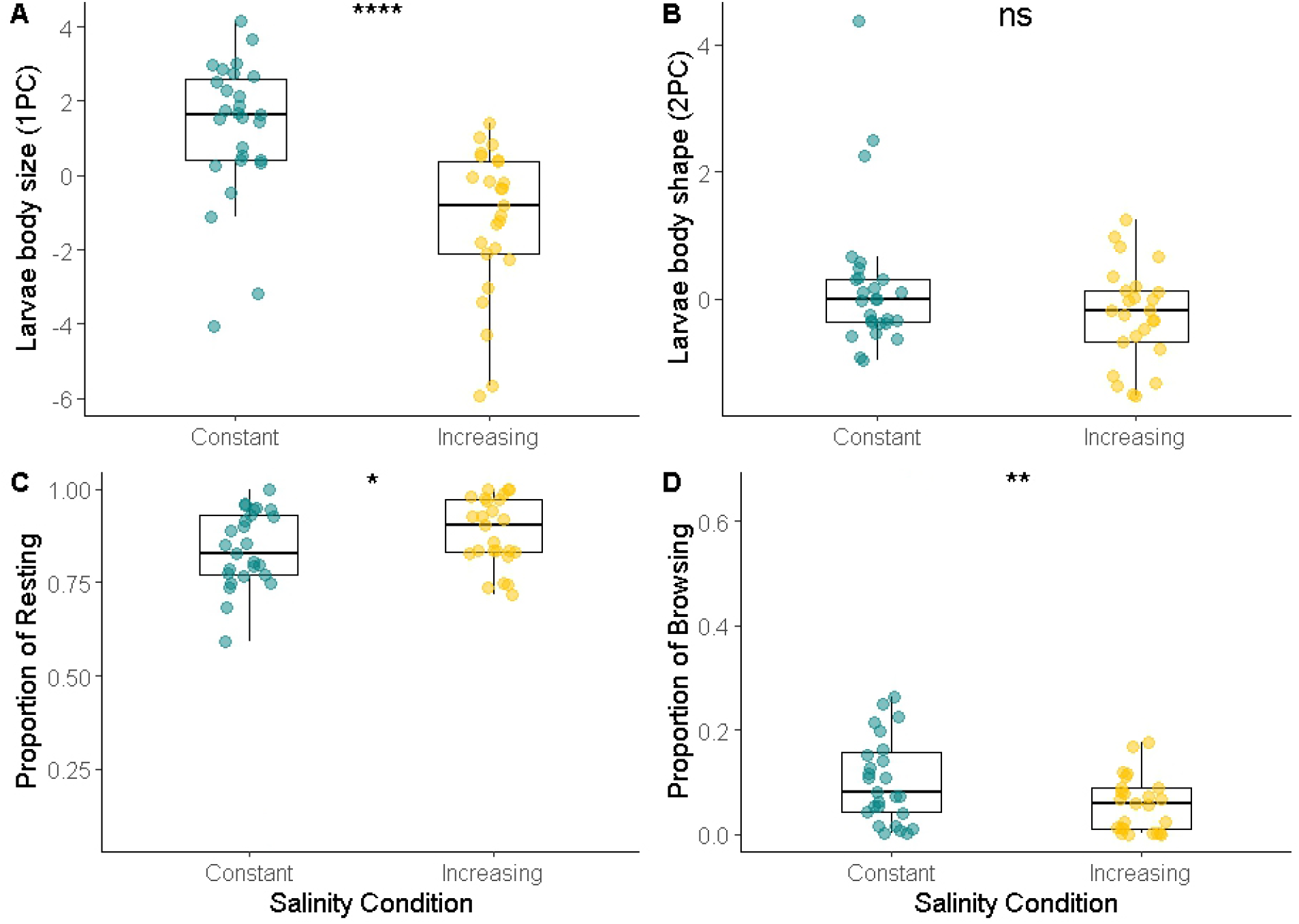
Larval phenotypic traits. A) PCA first component (larval size), B) PCA second component (larval shape), C) Proportion of time spent in resting and D) browsing of fourth instar larvae under constant (C; N = 27) and increasing salinity (S, N = 26) conditions. Points are individual observations. Significance levels: *** = *P* < 0.001, ** = *P* < 0.01, * = *P* < 0.05, ns = *P* > 0.05. Boxplots show median values (middle line), interquartile range (box), and range values, including some outliers (dots that extend beyond the min and max of the boxplot).

The activity behaviour of larvae varied significantly between treatments (Fig. 1C e D). Increasing salinity caused an increase in resting by 7.22% and reduced browsing behaviour by 6.31% compared to the constant treatment (Table 1).

### Path analysis

In constant salinity treatment, our path analysis revealed a strong positive impact of the larval size on the pupae cephalo-thorax (Fig. 3A and D). Resting behaviour did not influence pupae cephalo-thorax width (Fig. 3C and D) and didn’t depend on larval body size (Fig. 3B and D). Overall, bigger larvae result in bigger pupae.

**Figure 3.**
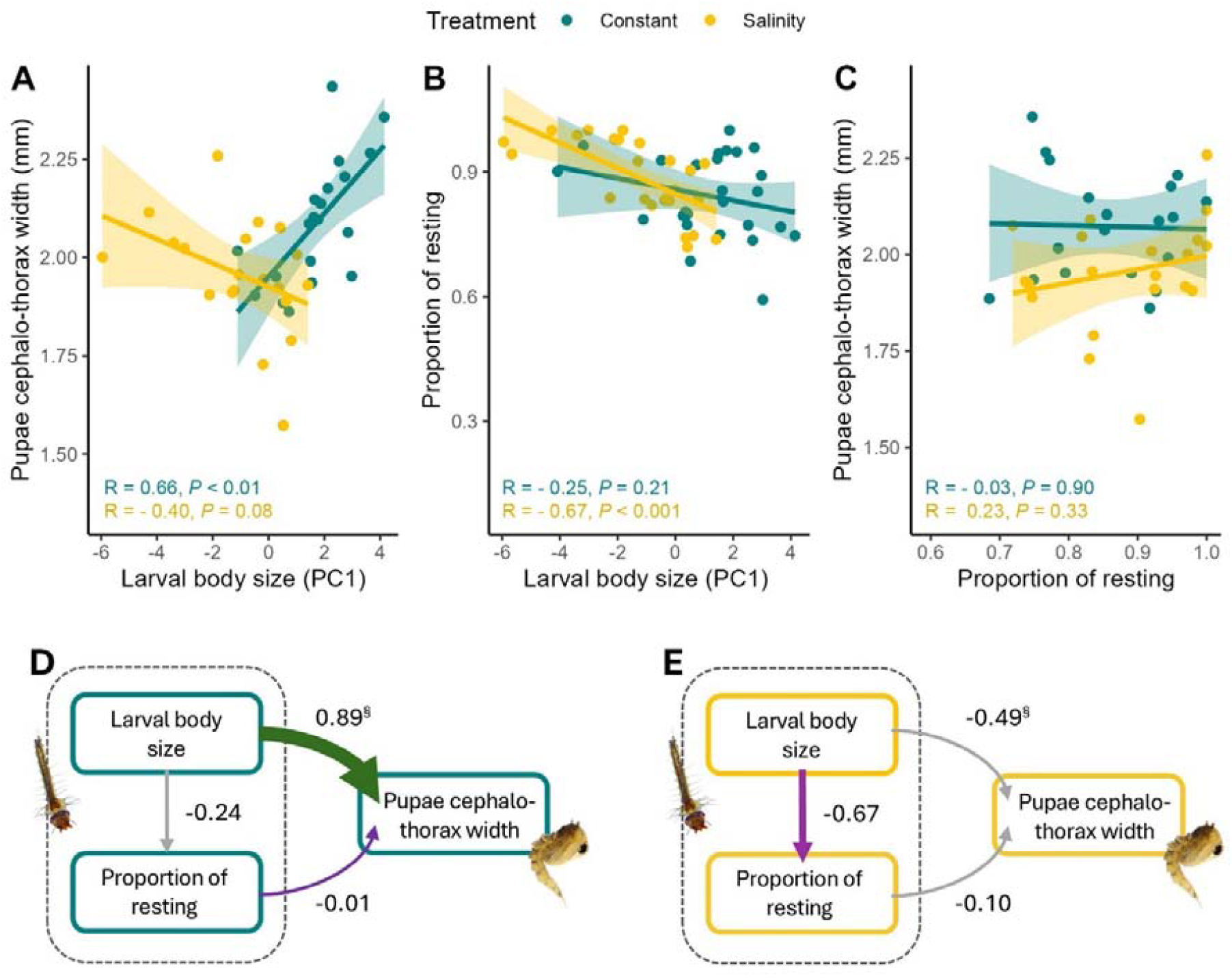
**A-C** Relationships between larval body size (PC1), pupal cephalothorax width, and resting proportion under two salinity conditions (Constant and Increasing Salinity). Each plot shows the linear regression relationship for the two conditions, with constant salinity represented by blue and increasing by yellow points and lines. Shaded regions represent the 95% confidence intervals for each regression line. Pearson’s correlation coefficient (R) and corresponding p-values are displayed for each condition within each panel. **D** Structural equation model (SEM) for constant treatment and **E** for increasing salinity treatment. Green and purple arrows represent significant (*P*<0.05) positive and negative paths, respectively, and grey arrows represent non-significant paths. Numbers next to the arrows are averaged effect sizes as standardised path coefficients; arrow widths reflect these standardised effect sizes. **§** represents paths with *P*□≤□0.05 in multigroup analysis (for exact *P*-values see Table 2). For the percentage of variance explained by response variables, non-standardised coefficient values and exact *P*-values of individual paths, see supplementary Table S3

Under increasing salinity conditions, the covariation patterns changed: cephalo-thorax width was no longer dependent on larval size (Fig. 3A and E). Moreover, in this salinity condition, the larval size negatively influenced the resting behaviour, meaning that smaller larvae perform more resting (and less browsing since there is a strong negative correlation between these two behaviours; Fig. 3B and E).

**Table 2.**
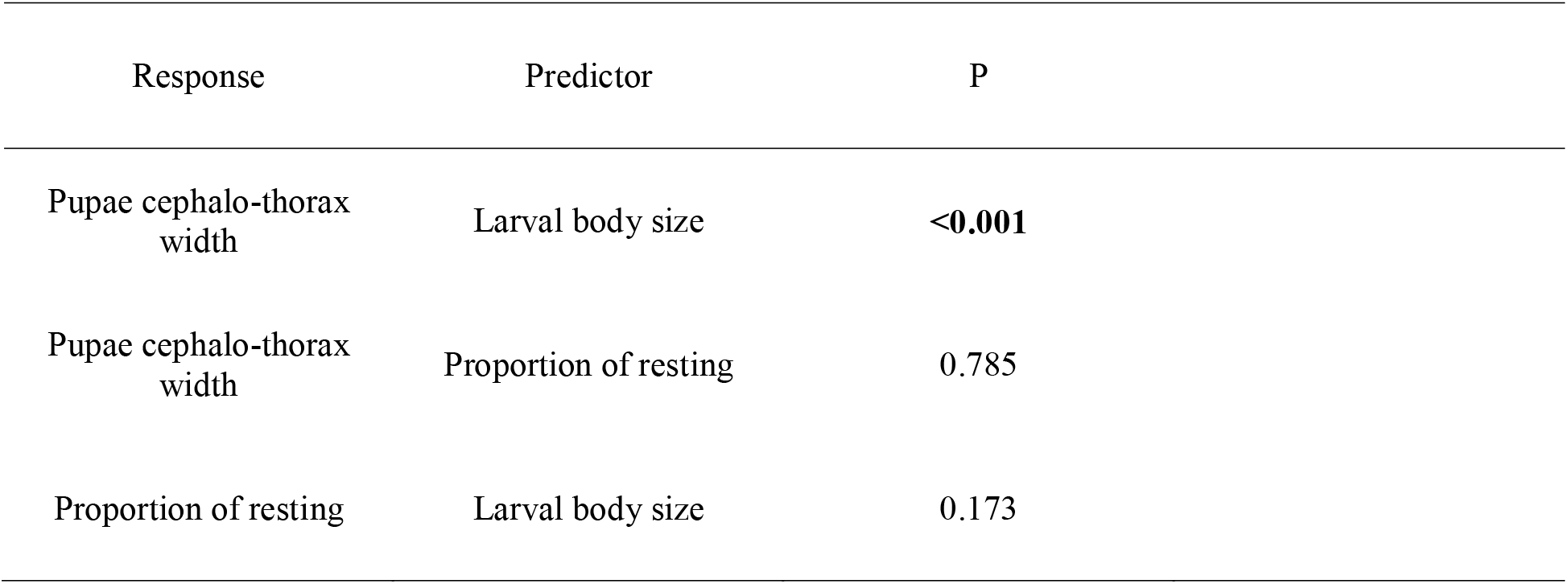
Exact P-values of the multigroup analysis. This analysis implements a model-wide interaction in which every term in the model interacts with the grouping variable (i.e. constant versus increasing salinity).

Finally, we refitted the SEM analysis using the whole dataset, including treatment as a factor. Results identified one statistically significant change in the pathways between treatments: the strong positive relationship between larval body size and pupae cephalo-thorax width became non-significant under increasing salinity (Fig.3D and E) (Table 2).

## Discussion

In this study, we measured larval activity and body size throughout metamorphosis to determine whether these traits exhibit plasticity under changing salinity conditions. We then performed a path analysis to infer causal relationships between body size and behavior during development and to test the hypothesis that such trait interactions can shape direct carry-over effects. Overall, our results show that salinity influences plastic changes in both morphology and behaviour of *Ae. mariae* larvae. Most importantly, we found that salinity affects trait covariation, which in turn influences body size development across metamorphosis, supporting our hypothesis.

First, we observed a plastic response to increasing salinity, resulting in smaller larval body size and reduced browsing behaviour (Fig. 1). These changes can be attributed to the energetic costs of osmoregulation. Several studies have shown that mosquito species inhabiting brackish or saline water maintain osmotic balance through different strategies, such as accumulating amino acids in the hemolymph [49,50], increasing the size of anal papillae [51] or producing less permeable cuticles [52]. These mechanisms require substantial energy expenditure, which is traded off against resource allocation for body growth [53–55]. Interestingly, our analysis of activity behaviour revealed that larvae in the increasing salinity treatment spent less time browsing (i.e., actively feeding) and more time resting compared to those in the constant salinity treatment. Since mosquito larvae filter and ingest water while resting at the surface, oral water intake in saltwater mosquito species may counteract osmotic water loss through the cuticle [45]. For instance, in *Culex tarsalis*, larvae transferred to higher salinity increased their drinking rate by over 50% compared to freshwater conditions [50]. Accordingly, the combined effect of energy allocation to osmoregulation and behavioral plasticity in *Ae. mariae* likely contributes to its smaller body size under increasing salinity conditions.

Secondly, using path analysis to examine the correlation between larval traits in the two treatments, we found no interaction between larval size and activity under constant salinity conditions. However, under increasing salinity, activity behaviour became size-dependent. Specifically, we observed a strong negative effect of larval body size on activity behaviour (Fig. 3B-E). This suggests that salinity conditions influence trait covariation. This finding aligns with existing literature, as it is well established that environmental conditions can shape not only the expression of individual traits but also their covariation, i.e., the plasticity of phenotypic integration [10,22,31,56]. For instance, Relyea (2001) found that in multiple species of larval anurans, the patterns of trait covariation between morphological and behavioural traits varied depending on predator exposure; In predator-free environments, activity was positively correlated with body depth and width, a pattern that was partially maintained (between activity and body width) in the presence of *Umbra* predators but entirely absent in the presence of *Anax* predators. Similarly, a negative correlation between activity and tail depth was observed in the absence of predators and in the presence of *Umbra* individuals but disappeared in the presence of *Anax* predators.

As we move from the larval to the pupal stage, we found that the plasticity of phenotypic integration influences phenotypic development throughout metamorphosis. Specifically, under constant salinity conditions—where no correlation between larval size and activity was observed— we detected a strong, positive, and significant effect of larval size on pupal size. This indicates that larger larvae developed into larger pupae, demonstrating a direct carry-over effect. Conversely, under increasing salinity conditions, the emergence of size-dependent activity behaviour completely disrupted the relationship between larval size and pupal size, effectively breaking the direct carry-over effect. Trait decoupling across metamorphosis can result from changes in genetic correlations between life stages or from differences in the environmental sensitivity of later-stage development [57–61]. However, in the case of direct carry-over effects, constraints on decoupling are typically strong, as later-stage phenotypes are inherently shaped by developmental factors acting on the same traits earlier in life [8,62]. Here, we provide evidence that interactions between plastic traits may serve as an alternative mechanism for decoupling in cases of direct carry-over.

## Conclusions

The extent to which one life stage is linked to others across metamorphosis remains a topic of debate. According to the ‘adaptive decoupling hypothesis’, the phenotype of one life stage does not influence other stages, allowing traits to evolve independently across developmental transitions [60,63]. Conversely, evidence of carry-over effects across a wide range of organisms with complex life cycles or those spanning multiple environments [9,17,64–66] suggests that metamorphosis does not represent an entirely new beginning, but rather a process in which different life stages remain developmentally connected. Our findings reveal that the connection between life stages across metamorphosis is not fixed but can be plastically reshaped by environmental stressors, leading to a ‘plastic’ direct carry-over effect.

## Acknowledgments

The authors thank Nicole Giardiello and Alessandra Spanò for their technical help.

## Notes

### Competing Interest Statement

The authors have declared no competing interest.

### Summary of Updates

New statistical analyses have been included focusing on larval stage. Accordingly, some parts of the text have been modified

